# The neural basis of creative production: A cross-modal ALE meta-analysis

**DOI:** 10.1101/2021.03.09.434575

**Authors:** Steven Brown, Eunseon Kim

**Affiliations:** Department of Psychology, Neuroscience & Behaviour, McMaster University, Hamilton, ON, Canada

**Keywords:** creativity, fMRI, improvisation, domain-specific, domain-general, divergent thinking, frontal aslant

## Abstract

One of the central questions about the cognitive neuroscience of creativity is the extent to which creativity depends on either domain-specific or domain-general mechanisms. To address this question, we carried out two parallel activation likelihood estimation meta-analyses of creativity: 1) a motoric analysis that combined studies across five domains of creative production (verbalizing, music, movement, writing, and drawing), and 2) an analysis of the Alternate Uses divergent-thinking task. All experiments contained a contrast between a creative task and a matched non-creative or less-creative task that controlled for the sensorimotor demands of task performance. The activation profiles of the two meta-analyses were non-overlapping, but both pointed to a domain-specific interpretation in which creative production is, at least in part, an enhancement of sensorimotor brain areas involved in non-creative production. The most concordant areas of activation in the motoric meta-analysis were high-level motor areas such as the pre-supplementary motor area and inferior frontal gyrus that interface motor planning and executive control, suggesting a means of uniting domain-specificity and -generality in creative production.

---

Creativity has garnered a great deal of attention in the psychology and neuroscience literatures in recent years (Abraham, 2018; Chan, 2015; Jung and Vartanian, 2018; Kaufman et al., 2017; Kaufman and Sternberg, 2019), in part because of its real-world application to understanding technology development, scientific discovery, and artistic creation, among many other domains. Articles in the popular press attempt to identify the traits and habits of creative people in order to provide strategies for increasing one’s own level of creativity (e.g., The Creativity Post website), since creativity is generally seen as being a valued personal attribute. However, there are many contentious issues regarding the nature of creativity (Weisberg, 2020, 2006), as will be described in the following sections. We shall begin with a theoretical review of key topics related to the nature of creativity and then examine some of these ideas empirically by performing an activation likelihood estimation (ALE) meta-analysis of neuroimaging studies of creativity, as described in the second half of the article.

## Creation and creativity

The term “creativity” is both easy to define and difficult to apply. In its most universal definition, creativity refers to the novelty of an idea or product, including its originality and level of surprise (Stein, 1953; Torrance, 1988). A non-creative idea or product is something that strongly resembles existing ideas or products, as related to concepts such as reproduction, replication, imitation, and re-use. Because humans are consummate conformists (Boyd and Henrich, 1998; Legare and Nielsen, 2015; Mesoudi and Lycett, 2009; Sternberg and Lubart, 1995), novelty is probably a deviation from the default state of human behavior, which is grounded in imitation, social learning, and an adherence to social norms (Boyd and Richerson, 2005, 1985) and historical traditions (Liénard and Boyer, 2006). However, there are many situations in which novelty can be socially valued, situations where people respond “constructively to existing or new situations, rather than merely adapting to them” (Torrance, 1988:47). In the realm of problem solving, creativity refers to the development of new strategies for solving a problem, rather than a reliance on existing solutions, especially when the latter do not work. In the realm of artistic production, it refers to the composition of a new work that did not previously exist, rather than the performance of a pre-composed work. Creativity is nothing if not a *relative* term, comparing a given idea or product to what has existed previously (Boden, 2010; Csikszentmihalyi, 1988; Stein, 1953).

Rhodes (1961) argued that the psychological study of creativity should be based on what are now called “the 3 P’s” of person, process, and product (Rhodes also has a 4^th^ P of press). Despite the utility of this trichotomy, it introduces significant problems in thinking about a definition of creativity. For example, the creativity of a person cannot be the same thing as the creativity of a product, and neither can be the same as the creativity of a cognitive process. Using the same word for all three concepts is problematic. Let us consider the three cases of person, product, and process, respectively (Figure 1).

**Figure 1.**
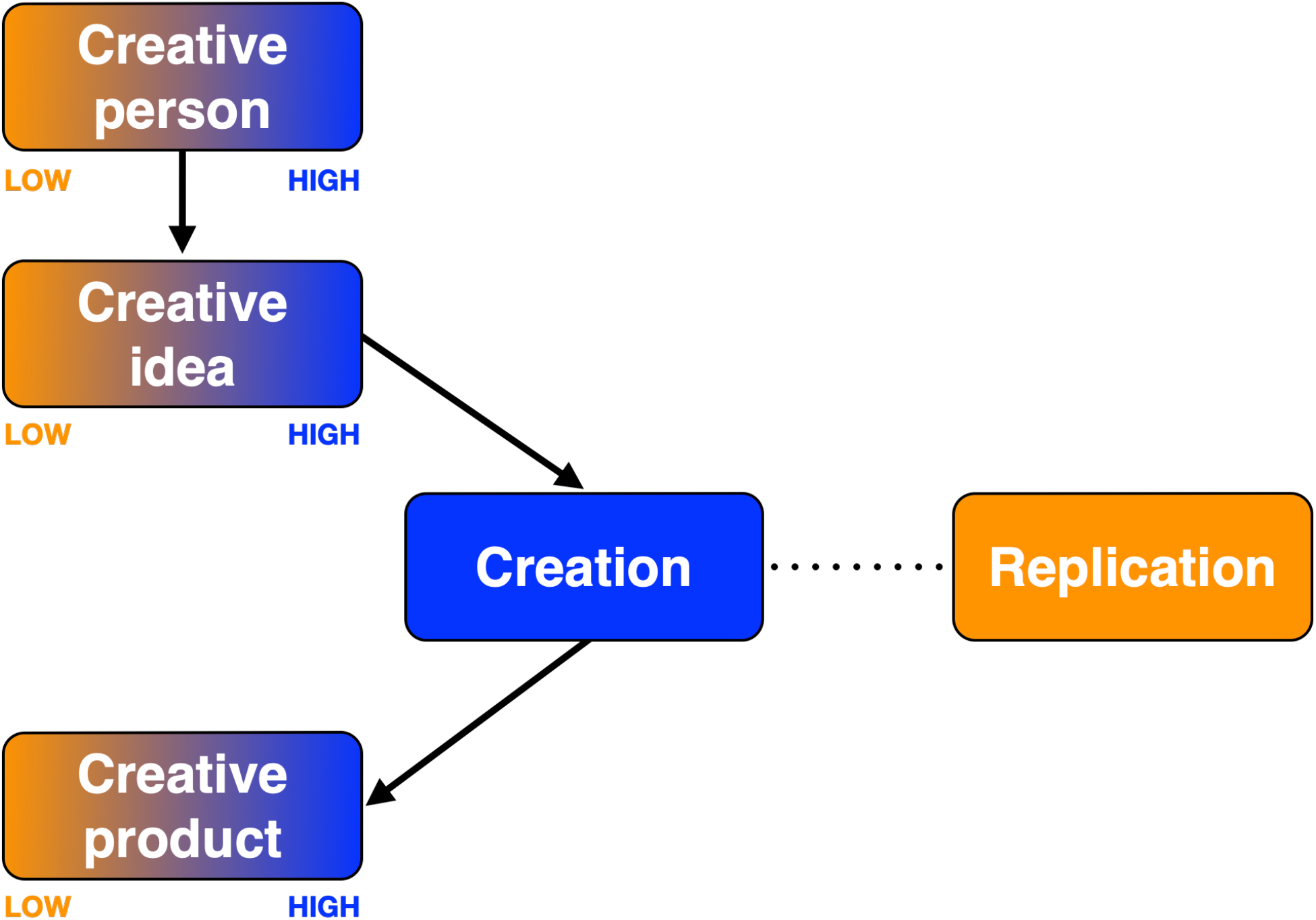
The graded nature of creativity compared with the categorical difference between creation and replication. Creativity is shown with shading spanning from low to high creativeness. Creation (i.e., the process creative production) is shown with a uniformly-colored box to contrast it with replication as non-creative production. Likewise, creation, as a categorical process, is shown in a separate column from the graded processes of creativity.

In talking about people as being more or less creative, one does not mean that they are more or less novel as human beings (where a term like “unique” might be more appropriate). What one means is that *their ideas* are more or less novel. Hence, when one talks about a person as being creative, what this really means is that the person is imaginative and that they have a strong capacity to generate original ideas, not least to solve problems in an innovative manner. The creativity of a person thus relates to this capacity to generate ideas, rather than to the ideas themselves. This capacity can be seen as both a trait – as in the study of the differential psychology of creativity – and a state, as in creators who have both fertile and dry periods across their careers. Historiometric analyses of creativity and eminence (Kozbelt, 2008; Martindale, 1995; Simonton, 2010, 1992, 1991) examine this capacity across an individual creator’s lifetime, and attempt to look for group trends across members of different domains of creative work. Psychological studies of personality (Feist, 2019), intelligence (Kim et al., 2010), mental illness (Carson, 2011; Simonton, 2014) and other trait-variables attempt to explain why some people are more imaginative than others when it comes to the generation of original ideas and products. However, the creativity of a person is about a capacity, not about the ideas that emanate from it. Creative people are simply imaginative individuals who generate creative ideas and products. The popular interest these days in “how to be more creative”, for example through creativity training programs, is not about learning how to become a free-spirited Bohemian, but about enhancing one’s capacity to generate novel ideas and products. The psychometric approach to creativity (Guilford, 1967; Plucker and Makel, 2010; Torrance, 1988) has been primarily driven by a search for ways to distinguish imaginative from non-imaginative people in the general population. Overall, the creativity-as-novelty concept does not apply to people per se, only to their ideas and products.

However, even ideas and products need to be distinguished from one another when it comes to creativity. For example, the popular “Four C” continuum for creativity (Kaufman and Beghetto, 2009; Simonton, 2010) – spanning from the everyday mini-C creativity of an inventive home chef to the domain-transforming Big-C creativity of an Einstein or Beethoven – applies to the *product* level, rather than the idea level. It refers to how the products of creativity impact a field of endeavour and drive its evolution. Similar ideas are found in Boden’s (2010) trichotomy of combinatorial creativity, exploratory creativity, and transformational creativity. The product level has social and historical connotations that the idea level does not (Csikszentmihalyi, 1988). For example, creative products are appraised by the members of a community as part of a process of cultural evolution to impact the transmission of those products across time and location. The idea level of creativity is different than this. It is often described as a style of “thinking”, as in the nearly ubiquitous assessment of creative cognition as divergent thinking in the cognitive psychology literature (Guilford, 1967; Runco, 2010; Torrance, 1988), itself contrasted with convergent thinking that is based analytically on antecedent ideas and products (Cropley, 2006; Guilford, 1967; Weisberg, 2020). For almost all fields, a creative idea is merely a potentiality if it is not translated into a product (“Wow, that’s a great idea”). Hence, it is only a starting point for an extended process of product generation.

What about the creative process? We propose replacing the process-related definition of creativity with the term “creation”, as shown in Figure 1. Creation is defined here as *the act of performing creative work*, regardless of the originality and the quality of the final outcome. It can occur in the moment as an improvisation, or it can extend over many years through long-term processes of collaborative work in large teams. Creation, as a production process, is placed in *categorical* distinction to replication, which is the non-creative process of imitatively reproducing something that already exists. Creation is the process of generating something novel, rather than employing an existing strategy or performing an existing work. It is the implementation of a creative idea to generate a product that can be shared publicly. The categorical nature of the distinction between creation and replication makes creation different from creativity, which refers to a *graded* appraisal of the level of novelty (originality) of something or someone, spanning from low to high creativeness. This is seen quite clearly in the Four C continuum for products and achievement. Hence, we propose distinguishing creation, as a categorical process distinct from replication and imitation, from the graded continua of creativeness that apply to the appraisals of the originality of people, ideas, and products. The caveat again is that there is a great need to develop qualifiers that distinguish the different senses of creativity that apply to people (imaginativeness), ideas (originality), and products (originality, but combined with the products’ cultural and historical impact). One important benefit of referring to the creative process as creation is that it allows us to move beyond the very idea-centric bias that is present in much research on the psychology of creativity and to appreciate the fact that creative work involves far more than simply generating an initial idea (Brown, 2019; Chan, 2015). Idea generation has acquired a privileged status over the processes that both precede and follow it in much theorizing about creativity. By contrast, the concept of creation includes *the full gamut of processes that extend from inspiration to final product*, with application to both improvisation and long-term processes of creative production.

While creation is placed in opposition to replication in Figure 1, the two processes are united by their requirement for a motor plan to carry out a goal-directed action and potentially to generate a product as well. Whether a pianist is playing Chopin or producing an improvisation, she needs to engage in the complex motoric act of playing the piano and, at a more cognitive level, of generating musical structure. This requirement for motor production and the communication of meaningful information is the same regardless of whether the performed product is novel or pre-composed. Because of this, creation is inextricably domain-specific, whereas there is still much debate about whether creativity, as an ideational process, is domain-specific or domain-general (see below). In addition, creation is strongly linked to mechanisms of *expertise*, whether this be motoric or cognitive. The jazz musician Charlie Parker was not only a highly innovative improviser but an extremely skilled saxophonist as well. Creation is often linked with problem solving in a given domain and to the implementation of functional solutions to problems (Weisberg, 2020), although the generation of novel products in the arts may occur for its own sake in the form of aesthetic displays, as seen in modernist trends in the arts (Goldberg, 2011). It should be pointed out that, in neuroimaging experiments of creativity, most parametric analyses of the creativeness of the generated products in the scanner use the creative-vs.-non-creative contrast in examining regressions with brain activity (e.g., Ellamil et al., 2012; Liu et al., 2015; Saggar et al., 2015). Hence, the analysis of creativity – as a graded feature of the generated products – is typically dependent on the analysis of creation as a categorical process distinct from replication.

## Combinatoriality and compositionality in creation

The distinction between creation and creativity is not just a semantic clarifier but an important consideration in developing theories about the evolutionary origins of creativity, cognition, and praxis. Many forms of creation in human cognition are based on assembling basic building-blocks into combinations. This is seen most notably in systems that are intrinsically combinatorial (generative) and compositional. Important examples include language and music. Jackendoff (2002) described how the three major systems of language processing – semantics, syntax, and phonology – are each generative systems with their own hierarchical combinatorial rules. Sentences are assembled through a compositional mechanism that combines words – as selected from a lexicon based on their meaning and their part of speech – to form syntactic constituents like noun phrases, which themselves combine to form clauses and phrases. Music has similar properties to the combinatoriality of phonology, whereby acoustic building-blocks are combined in a hierarchical fashion to form motives, melodies, and harmonies (Bernstein, 1976; Brown, 2017; Lerdahl, 2013; Patel, 2008). In visual art, “form primitives” such as geometric patterns serve as similar, though purely spatial, building blocks for visual composition in drawings and sculptures (Arnheim, 1974; Golomb, 2002).

Such observations have striking implications for the evolution of creativity. If human-specific systems like language and music are intrinsically based on combinatorial principles, then their functions in everyday interactions are inherently creational. During conversation, we are constantly improvising our utterances, even if their creative quality is not that of a master storyteller. Hence, the evolutionary leap for creative storytelling, for example, was the evolution of language and speech, not the evolution of some general creativity module. While the level of creativity of a given product may indeed be modulated by domain-general processes like working memory (Vartanian, 2019; Wynn and Coolidge, 2014), the very process of creation is linked to newly-evolved, domain-specific modules like language and music that are built on combinatorial operations at their core. Hence, we cannot talk about creativity in the abstract, nor can talk about it as a general “skill”. It has to be the creativity of something, and that something pertains to a particular domain of creation. For example, there could be no highly original music if humans didn’t first possess the combinatorial capacity to create music and to translate that capacity into products via vocalization and instrumental production. In the realm of the arts, creators (e.g., composers, choreographers, playwrights) are typically distinguished from performers. Creators engage in the act of creation, whereas performers engage in the act of reproduction, the major exception being improvisers, who engage in creation during the course of performance, much the way that all people improvise their utterances during conversation.

## Divergent thinking

“Divergent thinking” (DT) is the most popular operational definition of creativity in the psychology literature (Guilford, 1967). It is a type of outside-the-box thinking that focuses on the generation of novel ideas through brainstorming processes. The paradigms used to study it include a series of psychometric pencil-and-paper tests of creativity (Guilford, 1967; Kozbelt et al., 2010; Plucker et al., 2019; Torrance, 1988), most notably the Torrance Tests of Creative Thinking (Torrance, 1974). A standard means of examining DT in neuroimaging studies of creativity is to perform a comparison between the Alternate Uses task (AUT) – where people are asked to come up with unusual uses for common objects – and a control task that asks them to come up with ordinary uses for common objects (e.g., Abdul Hamid et al., 2019) or to describe the objects’ characteristics (e.g., Fink et al., 2010, 2009). While the participants’ responses can be registered through vocalization (e.g., Fink et al., 2009), a majority of studies have participants push a button to signal the occurrence of a new idea, and use this as a measure of the number of generated ideas (see Methods section). DT tasks like the AUT are purely ideational, and no true “product” is generated beyond a list of object uses. In addition, the task operates without concern for the utility of these uses (Baer, 2011; Perry-Smith and Mannucci, 2017), but simply their quantity and originality. Studies of DT, in contrast to those of motor improvisation, are typically carried out using non-specialist participants. From a cognitive standpoint, ideational tasks like the AUT are limited to a phase of idea generation, whereas most motoric tasks go beyond generation alone to engage in an elaboration of the ideas to create products, as in the production of a drawing, poem, or musical improvisation. The AUT, by contrast, does not have an elaboration phase.

## Domain and time-frame

A unified theory of creativity has to deal with two overarching dimensions of creative work: domain and time-frame (Figure 2). The *domain* dimension examines whether the mechanisms of creativity are cross-modal or are distinct for each domain. Are the mechanisms of piano improvisation the same as those for Tango dancing and improvisational acting? One of the aims of the meta-analysis described below is to examine creative production in a cross-modal manner by analyzing different domains of short-term creativity. The *time-frame* dimension of creativity distinguishes the improvisational type of creativity that occurs in the moment from the long-term creativity that takes place over extended periods of exploration and revision. Are the mechanisms of performing a piano improvisation the same as those involved in composing a piano sonata (Larson, 2005)? We will examine each of these dimensions in succession.

**Figure 2.**
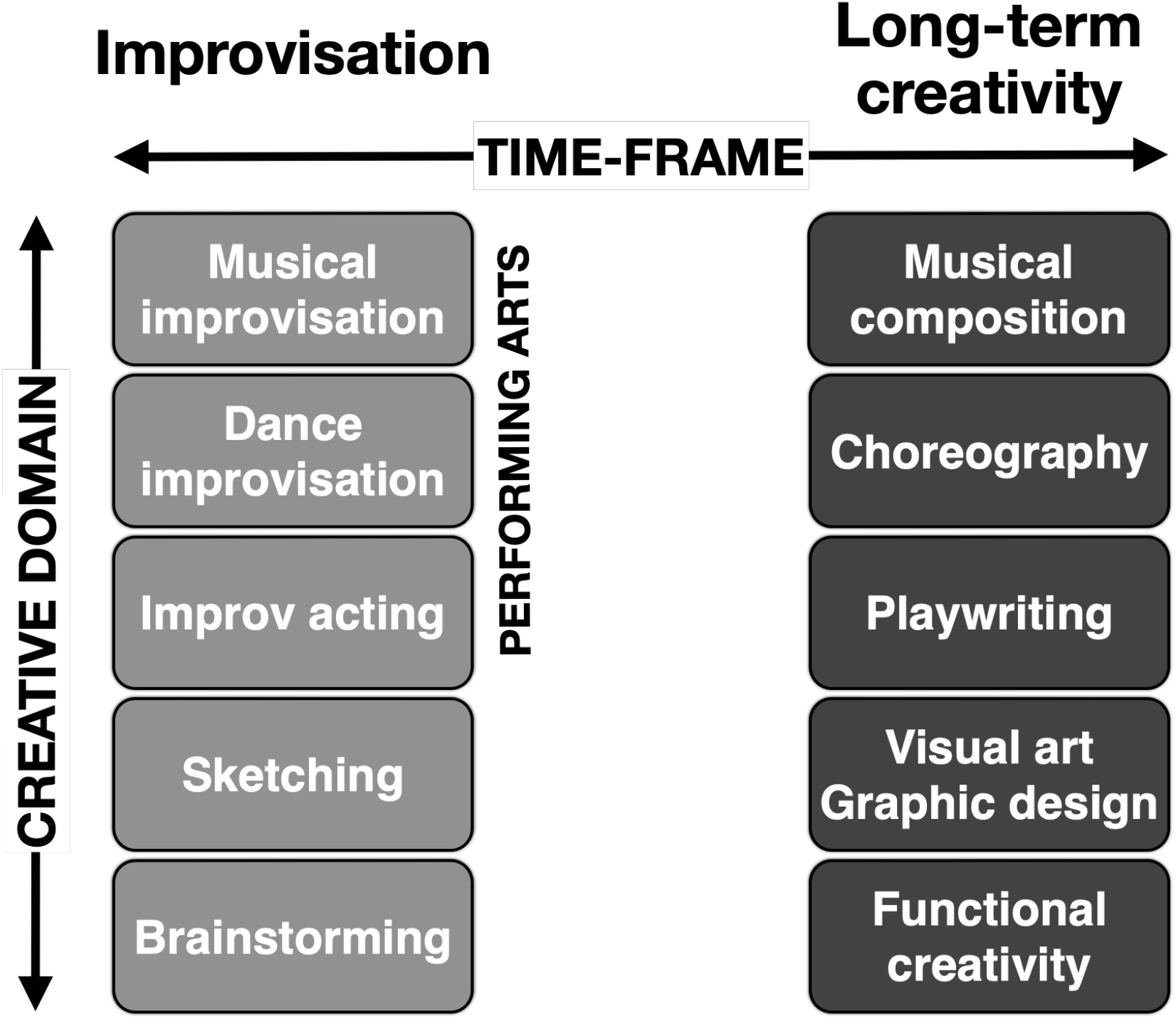
The domains and time-frames of creativity. This is shown using five example domains.

### Domain

One of the central questions about the cognition and neuroscience of creativity is the extent to which creativity depends on either domain-specific or domain-general mechanisms, or some combination of the two (reviewed in Baer, 2016; see also Boccia et al., 2015; Palmiero et al., 2010). Compelling arguments have been made on both sides of the debate (Baer, 2016, 2010; Plucker, 1998), suggesting that the answer is most likely some combination of the two types of mechanisms. Despite the widespread use of non-specialist participants in DT tasks in the lab, real-world creativity is carried out by specialists who work in a goal-directed manner to solve domain-specific problems (Amabile and Pratt, 2016; Brown, 2019; Glaveanu et al., 2013; Hennessey and Amabile, 1988; Weisberg, 2020). Such people have a “prepared mind” for carrying out creative work (Cropley, 2006). In order to shed light on the domain dimension of creativity from a neural standpoint, we will propose a distinction between the two neural mechanisms that we call *enhancement* and *expansion*. (Although there is no reason that these mechanisms need be dichotomized, we will present them in a dichotomous manner for the present purposes in order to highlight potential differences in the brain networks associated with each one.) Figure 3 presents a graphic model of these mechanisms using musical creativity as an example function.

**Figure 3.**
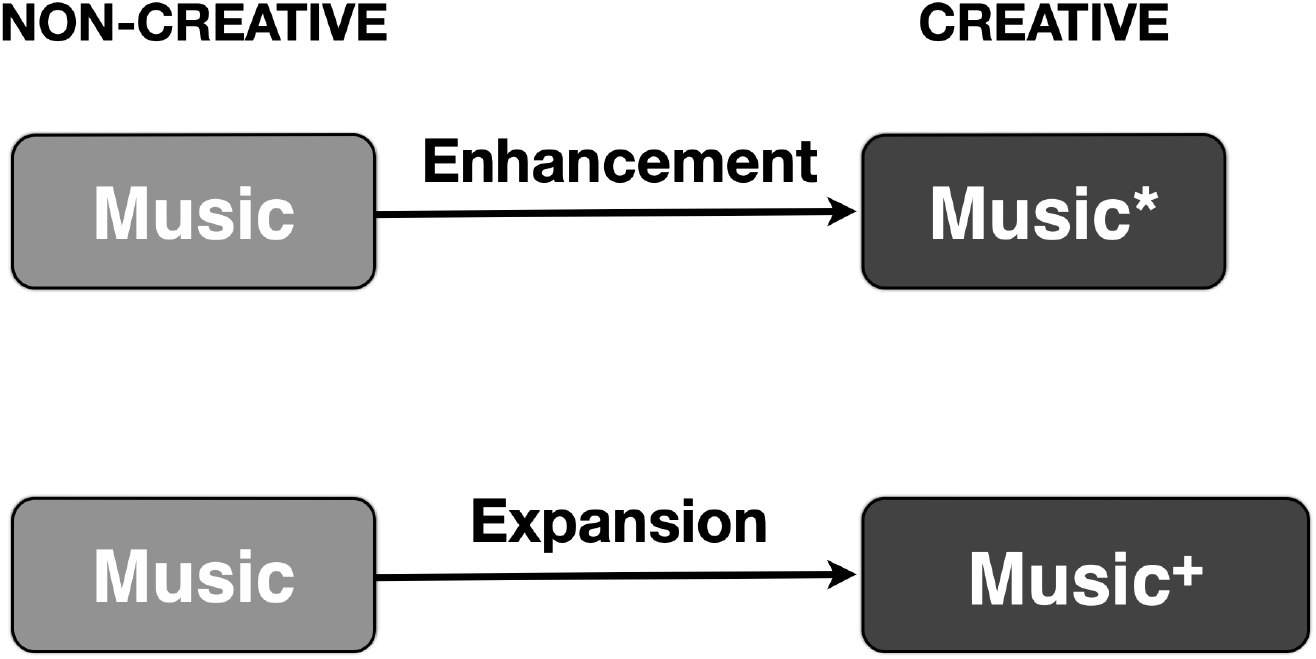
Two neural models of the transition from a non-creative to a creative state of processing. Music is used as an example function. Enhancement is a *domain-specific* mechanism in which the sensorimotor networks that mediate non-creative production are amplified during creative production, as indicated by the * symbol. Expansion is a *domain-general* mechanism in which processes like attention and working memory are expanded beyond their normal levels in the non-creative state to produce the creative state, as indicated both by the + symbol and the expansion of the box size.

Enhancement is conceptualized as a *domain*-*specific* mechanism of transitioning from a typical, less-creative state of cognitive processing to a more-creative state by means of an activation increase in the same motor, sensory, and/or sensorimotor areas that mediate non-creative production. Expansion, by contrast, is conceptualized as a *domain*-*general* mechanism of broadening cognitive-processing capacity, for example through increases in attention, working memory, or associative processing. Potential neural correlates of expansion might include increased activity in brain areas controlling working memory (e.g., dorsolateral prefrontal cortex), attention (e.g., anterior cingulate cortex), the default mode of activity (the default mode network [DMN]), or associative memory (e.g., hippocampus, parahippocampal gyrus, anterior temporal lobe). Perhaps the most discussed mechanism of expansion in the creativity literature is “frontal disinhibition” (Carson, 2014; Jung and Haier, 2013) or the release of inhibitory top-down executive control so as to foster an openness to novel ideas and associations.

### Time-frame

While many texts acknowledge the central importance of domains to the study of creativity, very few consider the issue of time scales. Creative production can occur in the moment as a spontaneous activity (e.g., jazz improvisation, conversational discourse) or, more generally, it can occur over long periods of development, such as the multi-year creation of a new art work, technological product, or educational curriculum (see Figure 2). Not surprisingly, psychological studies of creativity in the lab have favored the short time scale of generation, and neuroscientific and genetic studies of creativity have typically adopted their methods. However, most of the real-world production of creative products occurs over a long time period, and therefore much of the psychology literature on creativity fails to do justice to the ecological validity of long time scales, instead emphasizing rapid tasks that can be done in the psychology lab or MRI scanner. As mentioned above, DT tasks like the AUT place their focus on idea generation alone. However, the vast majority of creative work is not limited to a brainstorming phase alone but includes extensive processes of elaboration, exploration, and revision (Chusilp and Jin, 2006; Lonergan et al., 2004; Mace and Ward, 2002; Perry-Smith and Mannucci, 2017), much as occurred in the writing of this article. Therefore, generation alone does not provide a sufficient picture of the creative process. The AUT is one of the very few approaches to the study of creativity that relies on a generative phase alone without including elaboration or exploration. In this regard, this task, despite its dominant status in the field (Benedek et al., 2019), is an outlier compared to real-world creative tasks.

An influential formalization of the creative process can be found in Finke, Ward and Smith’s Geneplore model (Finke et al., 1992; Mace and Ward, 2002; Ward and Kolomyts, 2019, 2010), in which creative production proceeds through iterative cycles of the *gene*ration and ex*plor*ation of ideas. Similar proposals have been made for collaborative creativity (Ness and Dysthe, 2020). The starting point for creative thinking is a germinal idea (Bennett, 1976). This idea is neither random nor fully formed (Locher, 2010; Mace and Ward, 2002), but is instead the starting point for a long process of exploratory work in order to bring it closer to a desired outcome. The exploration phase will often lead to the generation of new ideas along the way, making the creative process inherently iterative, elaborative and revision-based (Calic et al., 2020; Finke et al., 1992; Lonergan et al., 2004; Mumford et al., 2012; Perry-Smith and Mannucci, 2017). As a result, creative work can be modelled using a genealogy tree to track the evolution of ideas/products over a creator’s work time (Chan and Schunn, 2015; Chusilp and Jin, 2006; Weisberg, 2004). In the best case, an improved version of the germinal idea will emerge after multiple rounds of generation and exploration, taking place over the course of weeks, months or even years. In the worst case, the idea might prove to be flawed or untenable and will be abandoned (Mace and Ward, 2002).

Whether the creative process is a Darwinian mechanism is a matter of debate (Campbell, 1960; Gabora, 2013, 2005; Simonton, 1999), but there is no question that creative exploration is a highly *selective* process (Simonton, 1999) in which the best solutions move forward and the worst ones are filtered out through a process of optimization, just as in general problem solving (Weisberg, 2020) and operant conditioning (Stahlman et al., 2013). Because creative work generally produces a set of interim sketches and products, a better metaphor for creative work than biological evolution (Campbell, 1960; Simonton, 1999) is *cultural evolution*. Creative work proceeds at the local level of the creative group in a very similar manner to cultural evolution at the culture-wide level, resulting in a progressive change in the stylistic features of the product over time. Of course, once a creative product is completed and enters the culture at large, it then undergoes the normal process of cultural evolution at the culture-wide level.

While intuitive thinking about creativity sees the creative process as an act of problem solving (Guilford, 1967), an underappreciated aspect of creativity is problem *finding* (Abdulla et al., 2020; Guilford, 1950; Runco, 1994), in other words a determination of which problems within a given domain are worthy of being addressed to begin with. Sternberg (1988), in talking about the renowned experiments in the history of psychology, argued that “[p]roblem selection was far more important than problem solution in determining the classic role of these experiments” (p. 133). A century earlier, Souriau (1881) pointed out that “the truly original mind is that which discovers problems” (quoted in Campbell, 1960:385). Most creative work involves addressing a particular problem in a given domain, in which case the analysis of the problem is typically the first step in the creative process (Chusilp and Jin, 2006). Problem finding reveals a highly domain-specific aspect of creative work, since problem discovery is generally linked to a particular domain.

## Objectives of the meta-analysis study

Having presented a theoretical discussion of general issues related to the nature of creativity, we will now proceed to examine some of these ideas through a quantitative meta-analysis of neuroimaging studies of short-term creative processing; neuroimaging studies of long-term, explorational creativity have yet to be carried out. We will do this by performing two parallel meta-analyses of neuroimaging studies of creativity. One is a study of divergent thinking using the AUT. This should place the focus on generative brainstorming processes. The other is an analysis of motoric improvisation using all of the domains for which published studies are available. This should look at processes that incorporate elaboration in addition to generation in producing creative products, as in jazz improvisation and creative drawing. This analysis will cover five domains of improvisation: verbalization, music, movement, writing, and drawing, as shown in Figure 4. However, we are unable to compare these domains among themselves. Rules of best-practice for ALE meta-analysis require that there be no fewer than 17 matched experiments in order to carry out a robust analysis and obtain valid results (Eickhoff et al., 2016; Müller et al., 2018). Given that none of the five domains achieved this threshold, we combined all of the domains into a single “motoric” meta-analysis of 21 experiments (see Figure 4) and then compared this with 16 AUT experiments using a conjunction analysis. Note that this does not preclude an identification of cross-modal effects in the motoric analysis, since ALE clusters can be examined with respect to the domains that contribute to them. If a given ALE cluster is reliably found as a significant activation in the primary publications across the five domains of production, then this would argue for the cross-modal engagement of that brain region, even though the current state of the literature prevents us from running independent ALE analyses for each domain (although see Chen et al., 2020). An important aim of the meta-analyses is to examine whether the ALE clusters in each analysis correspond with either domain-specific or domain-general brain areas, as described above, in order to see if creativity occurs through mechanisms of enhancement, expansion, or both. Given that many neural models of creativity focus on domain-general networks like the DMN as the source of creative ideation (Beaty et al., 2016, 2015; Jung et al., 2013), then a domain-general model was our default prediction for both meta-analyses, especially for the AUT analysis.

**Figure 4.**
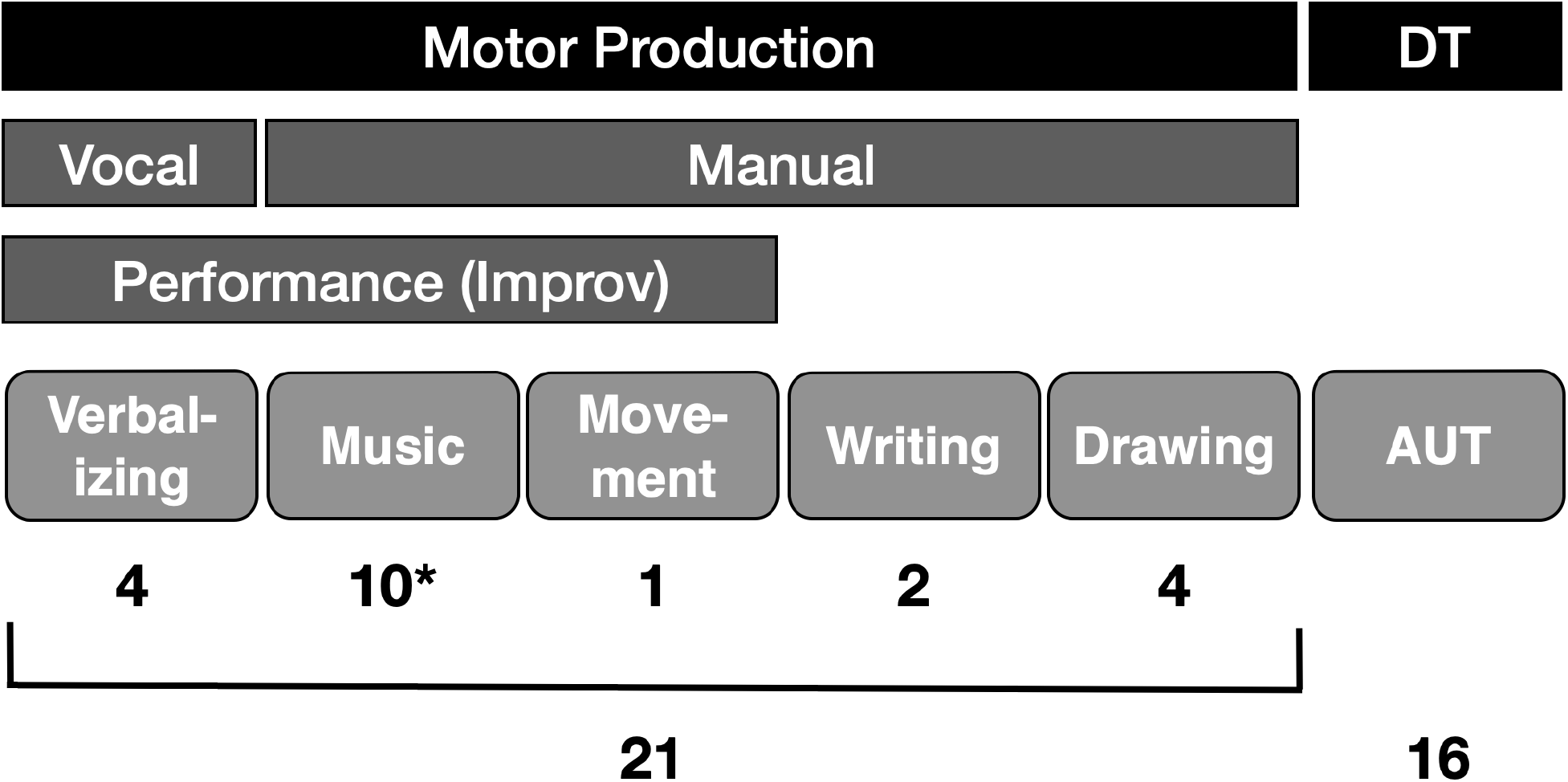
A categorization of the studies included in the meta-analyses into six categories. The number of experiments is shown below each box. The asterisk (*) for Music indicates that, whereas nine of the included studies are manual, one study is vocal. Abbreviations: AUT, Alternate Uses task; DT, divergent thinking.

The current approach to meta-analyzing the fMRI literature on creativity differs from previous meta-analyses in three manners. First, we restricted our inclusion of studies to those containing a contrast between a creative task and a matched non-creative or less-creative control task in order to equate the sensorimotor demands of task performance. Hence, we excluded studies that used rest or fixation as the baseline conditions, although most of the previous meta-analyses have (validly) included such studies as well (Boccia et al., 2015; Q. Chen et al., 2020; Gonen-Yaacovi et al., 2013). If sensorimotor areas are present in the matched contrasts, it would be indicative of a creativity-driven effect, rather than a task-related effect. Hence, this approach allows us to examine a sensorimotor model of creativity. Second, we chose to restrict the divergent-thinking meta-analysis to the AUT alone, while previous meta-analyses have included a broader coverage of tasks, including verb generation, visual imagery, metaphor production, and creative drawing (Cogdell-Brooke et al., 2020; Wu et al., 2015). Restricting the analysis to a single task would allow us to examine the reliability of activations for a single paradigm, while still covering the most universal divergent-thinking task in both the cognitive and neural literatures (Benedek et al., 2019). Third, it should be noted that different meta-analyses employ different schemes for organizing the published literature into functional categories. For example, both Gonen-Yaacovi et al. (2013) and Boccia et al. (2015) have a “verbal” category that includes not only verbal DT tasks like the AUT but also other verbal tasks like creative writing that we have placed in the “motoric” category in our analysis. Such differences need to be borne in mind when comparing the various meta-analyses among themselves.

## Methods

Activation likelihood estimation (ALE) meta-analysis is a coordinate-based statistical method to look for concordant areas of activation across a set of neuroimaging studies (Turkeltaub et al., 2002). Each focus of activation is modeled as a three-dimensional Gaussian probability distribution whose width is determined by the size of the subject group so as to reflect increasing certainty with increasing sample size (Eickhoff et al., 2009). Maps of activation likelihoods are created for each study by taking the maximum probability of activation at each voxel. A random-effects analysis tests for the convergence of activations across studies against a null hypothesis of spatially independent brain activations.

### Search query and inclusion criteria

We searched the PubMed database for published functional magnetic resonance imaging (fMRI) and positron emission tomography (PET) studies using the search terms “creativity”, “improvisation”, “figural creativity”, “alternate uses”, “creative writing”, “verbal creativity” and “music”. The reference sections of the retrieved publications were searched for additional studies. We classified the papers into six categories based on the type of creative task used in the experiment (see Figure 4). Five were motoric tasks (verbal improvisation, musical improvisation, movement improvisation, creative writing, and creative drawing), and one was an ideational divergent-thinking task, the AUT. Studies examining insight during problem solving were excluded (see Shen et al., 2018 for an ALE meta-analysis). A complete list of published studies containing included experiments can be found in Supplementary Table 1. Two separate ALE analyses were run: 1) the combination of the motor improvisation tasks, and 2) the AUT. Note that the meta-analyses only examined activation *increases* in the relevant subtractions. Too few articles reported deactivations to permit us to meta-analyze creativity-related deactivations. Individual articles, such as Limb and Braun (2008) and Liu et al. (2012), discuss such effects and their implications for the neuroscience of creativity.

**Table 1.** Talairach coordinates of the ALE clusters for the motoric and AUT meta-analyses. Stereotaxic coordinates are presented in millimeters along the left-right (x), anterior-posterior (y), and superior-inferior (z) axes. The “ALE” column provides the ALE score for each focus. Abbreviations: BA, Brodmann area; IFG, inferior frontal gyrus; PCC, posterior cingulate cortex; pre-SMA, pre-supplementary motor area; SMG, supramarginal gyrus.

|  | Motoric Tasks |  |  |  |  | Alternate Uses Task (AUT) |  |  |  |
| --- | --- | --- | --- | --- | --- | --- | --- | --- | --- |
|  | BA | x | y | z | ALE | x | y | z | ALE |
| FRONTAL |  |  |  |  |  |  |  |  |  |
| Pre-SMA | 6 | -6 | 10 | 48 | 0.034 |  |  |  |  |
|  |  | -8 | 6 | 60 | 0.023 |  |  |  |  |
| Frontal operculum | 44/45 | 44 | 22 | 10 | 0.023 |  |  |  |  |
|  |  | 46 | 12 | 6 | 0.021 |  |  |  |  |
|  |  | 48 | 10 | 16 | 0.020 |  |  |  |  |
| Dorsal IFG | 44 | 42 | 2 | 28 | 0.020 |  |  |  |  |
|  |  | -50 | 6 | 20 | 0.025 |  |  |  |  |
|  |  | -48 | 0 | 28 | 0.021 |  |  |  |  |
| Premotor cortex | 6 | -24 | -14 | 50 | 0.024 |  |  |  |  |
|  |  | -28 | -4 | 58 | 0.021 |  |  |  |  |
| Frontal operculum | 45 | -46 | 26 | 10 | 0.020 |  |  |  |  |
|  |  | -44 | 16 | 4 | 0.014 |  |  |  |  |
| TEMPORAL |  |  |  |  |  |  |  |  |  |
| Fusiform gyrus | 37 | -46 | -52 | -12 | 0.019 |  |  |  |  |
|  |  | -48 | -58 | -18 | 0.018 |  |  |  |  |
| PARIETAL |  |  |  |  |  |  |  |  |  |
| Anterior SMG | 40 |  |  |  |  | -54 | -32 | 36 | 0.029 |
| PCC | 23 |  |  |  |  | -14 | -56 | 20 | 0.018 |
| PCC | 31 |  |  |  |  | -8 | -58 | 26 | 0.012 |
| OCCIPITAL |  |  |  |  |  |  |  |  |  |
| Visual cortex | 17 |  |  |  |  | 10 | -84 | 8 | 0.024 |
| INSULA |  |  |  |  |  |  |  |  |  |
| Posterior insula |  |  |  |  |  | -34 | -10 | 2 | 0.021 |

The inclusion criteria for the studies in the meta-analyses were as follows: 1) that functional brain scanning was performed using either fMRI or PET, thereby excluding studies using electroencephalography, magnetoencephalography, functional near infrared spectroscopy, structural imaging techniques, and resting-state functional connectivity; 2) that the papers reported activation foci in the form of standardized stereotaxic coordinates in either Talairach space or Montreal Neurological Institute (MNI) space; 3) that results from the entire scanned brain volume were reported, thereby excluding studies that had partial brain coverage, that reported activation data for only specific areas, or that only reported region-of-interest analyses (e.g., de Manzano and Ullén, 2012a), with the exception of Limb and Braun (2008), who used a masking procedure; 4) that the participants were healthy adults, thereby excluding studies using clinical populations and healthy non-adults; 5) that the study contains a contrast between a creative task and a matched non-creative (i.e., replicative) or less-creative control task, thereby excluding studies using rest or fixation as the baseline conditions (e.g., Benedek et al., 2014) or that compared two creative conditions between themselves (e.g., Perchtold et al., 2018; Pinho et al., 2016), although an exception was made for Ellamil et al. (2012), who reported a create > evaluate contrast; 6) that the participants performed a generative task, thereby excluding studies in which participants evaluated other people’s work as creative (e.g., Mayseless et al., 2014); and 7) that a standard subtraction analysis was reported, thereby excluding studies that only reported a conjunction analysis, correlation analysis, connectivity analysis, or multivoxel pattern analysis (e.g., Beaty et al., 2015; Gilbert et al., 2010; Matheson et al., 2017; Pinho et al., 2016, 2014; Rutter et al., 2012), with the exception of de Manzano and Ullén (2012), who reported their results in a workable form using a conjunction analysis. The complete list of contributing experiments can be found in Supplementary Table 2.

Because 10 of the 16 AUT studies used a covert task with no vocal production, we permitted the inclusion of covert motor tasks in the motoric analysis as well (e.g., Amir et al., 2016; Hahm et al., 2017). For the graphical tasks, we only included creative drawing tasks (either overt or covert) and excluded tasks that involved graphic design but without a true drawing component. Hence, tasks that simply called for the assembly of objects from presented parts were excluded (e.g., Alexiou et al., 2011; Aziz-Zadeh et al., 2013; Cai et al., 2018; Goel and Vartanian, 2005). Beeson et al. (2003) was excluded from the sample of creative writing since the response only involved the production of single words, while the rest of the motoric studies included the production of full-fledged phrases, like poems, raps, and musical melodies. For this reason, we also excluded studies of verb generation (e.g., Seger et al., 2000), although other meta-analyses have included them.

Some articles that met our inclusion criteria reported two or more very similar contrasts that differed only slightly in emphasis, generally with nearly identical activation profiles. We refer to these as “duplicate” experiments from the standpoint of meta-analysis. For example, in Berkowitz and Ansari’s (2008) study of piano improvisation, one contrast consisted of melody improvisation vs. playing pre-learned melodies, while another contrast consisted of rhythm improvisation vs. improvising using a fixed rhythm. The results were very similar. In situations such as this, we aimed to be conservative by excluding potentially duplicate results from a single article, since retaining them would have artificially increased the concordance of the activated regions (Müller et al., 2018). The experiment that was selected from the two or more closely related experiments was the one that best matched the tasks in the other studies of that category, without any consideration for the results themselves. This impacted the following articles that met our inclusion criteria: 1) in the Verbalizing category, Bechtereva et al. (2004); 2) in the Music category, Berkowitz and Ansari (2008), Villarreal et al. (2013), Donnay et al. (2014), McPherson et al. (2016), and de Aquino et al. (2019); and 3) in the AUT category, Benedek et al. (2018), Abdul Hamid et al. (2019), and Madore et al. (2019). The excluded duplicate experiments from these articles are indicated in a separate column in Supplementary Table 2. In contrast to this situation, Fink et al. (2015) ran two separate participant groups in their training study of DT using the AUT. While the participants were tested on three separate occasions, the initial time point, called T1, was a common baseline for both training groups. Hence, we included the T1 data for the two training groups as two separate experiments. This is the only publication for which two independent experiments are included for either of the meta-analyses.

The meta-analyses included 37 experiments (421 foci, 857 participants) from 36 published studies, as organized into 21 experiments for the motoric meta-analysis (318 foci, 408 participants) and 16 experiments for the AUT analysis (103 foci, 449 participants). This is schematized in Figure 4. The 21 studies for the motoric analysis are comprised of 4 vocal experiments and 17 studies of manual production. These latter included 10 experiments of musical improvisation (128 foci), one of movement improvisation (16 foci), two of creative writing (48 foci), and four of creating drawing (92 foci). Note that one of the musical studies is actually vocal (Dhakal et al., 2019), but it is still listed in the Manual/Music category.

All analyses were performed using GingerALE 3.0.2 (www.brainmap.org/ale) according to standard methods (Eickhoff et al., 2016, 2012, 2009; Müller et al., 2018). MNI coordinates were converted to Talairach coordinates within GingerALE. The meta-analyses were performed as 5000 threshold permutations using a cluster-level, family-wise error threshold of p<0.05 and a cluster-forming threshold of p<0.001. The ALE scores in Table 1 are a reflection of the effect sizes reported in standard meta-analyses outside of the neuroimaging field (Eickhoff et al., 2012). The ALE results were registered onto a Talairach-normalized template brain using Mango 4.1 (ric.uthscsa.edu/mango).

In order to examine the reliability of the results in the published literature, we performed a diagnostic “contribution analysis” (CA) that classifies the activations reported in the coordinate tables of the published articles with regard to five functional categories of brain areas: 1) motor areas (premotor cortex, supplementary motor area [SMA], cerebellum, and basal ganglia); 2) pre-SMA and inferior frontal gyrus (pre-SMA, dorsal Brodmann area [BA] 44, and the frontal operculum [both BA 44 and 45]); 3) sensory cortex (e.g., visual cortex, auditory cortex) and thalamus; 4) domain-general executive control areas (dorsolateral prefrontal cortex [DLPFC] and anterior cingulate cortex [ACC]); and 5) the default mode network (posterior cingulate cortex [PCC], temporoparietal junction [TPJ], medial prefrontal cortex [mPFC], and anterior temporopolar cortex [ATPC]). Note that many of the areas described in the CA *do not* correspond with ALE clusters in the meta-analyses since activations in these regions were not sufficiently concordant across the experiments in the meta-analyses to form significant ALE clusters at the thresholds used. However, these areas *are* statistically significant at the level of the individual experiments. Hence, it is legitimate to report what percentage of the papers in the meta-analyses report activations in these five functional categories of brain areas as a measurement of reliability across the studies, just as in meta-analyses in other domains.

## Results

Figure 5 presents the results for the two meta-analyses: the motoric ALE (21 experiments across 5 domains of production) and the AUT ALE (16 experiments in a single domain). The Talairach coordinates of the ALE foci are listed in Table 1, and the results of the contribution analysis are shown in Table 2. Figure 5a shows the ALE foci for the motoric meta-analysis. The strongest foci occurred in the left pre-SMA, the dorsal part of BA 44 bilaterally, and the frontal operculum bilaterally (BA 44/45). Based on the CA, this group of areas was present in 86% of the motoric experiments (see Table 2). Next in importance were the premotor cortex (BA 6) and the visual association cortex of the fusiform gyrus (BA 37). In general, motor and sensory areas were each present in 67% of the contributing studies. Next, the CA shows that the DLPFC and/or ACC were present in 67% of the motoric experiments, although the activation coordinates for these areas were not sufficiently overlapping to produce any ALE clusters at the current threshold. Components of the DMN were present in 24% of the studies, most commonly the medial prefrontal cortex in BA 10. The PCC was present in only one of the 21 experiments (i.e., the creative writing study of Shah et al., 2013). The CA for the motoric meta-analysis shows an overall pattern of reliability since all of the ALE clusters have strong contributions from the component studies, including from all five of the motor domains in the meta-analysis.

**Table 2:**
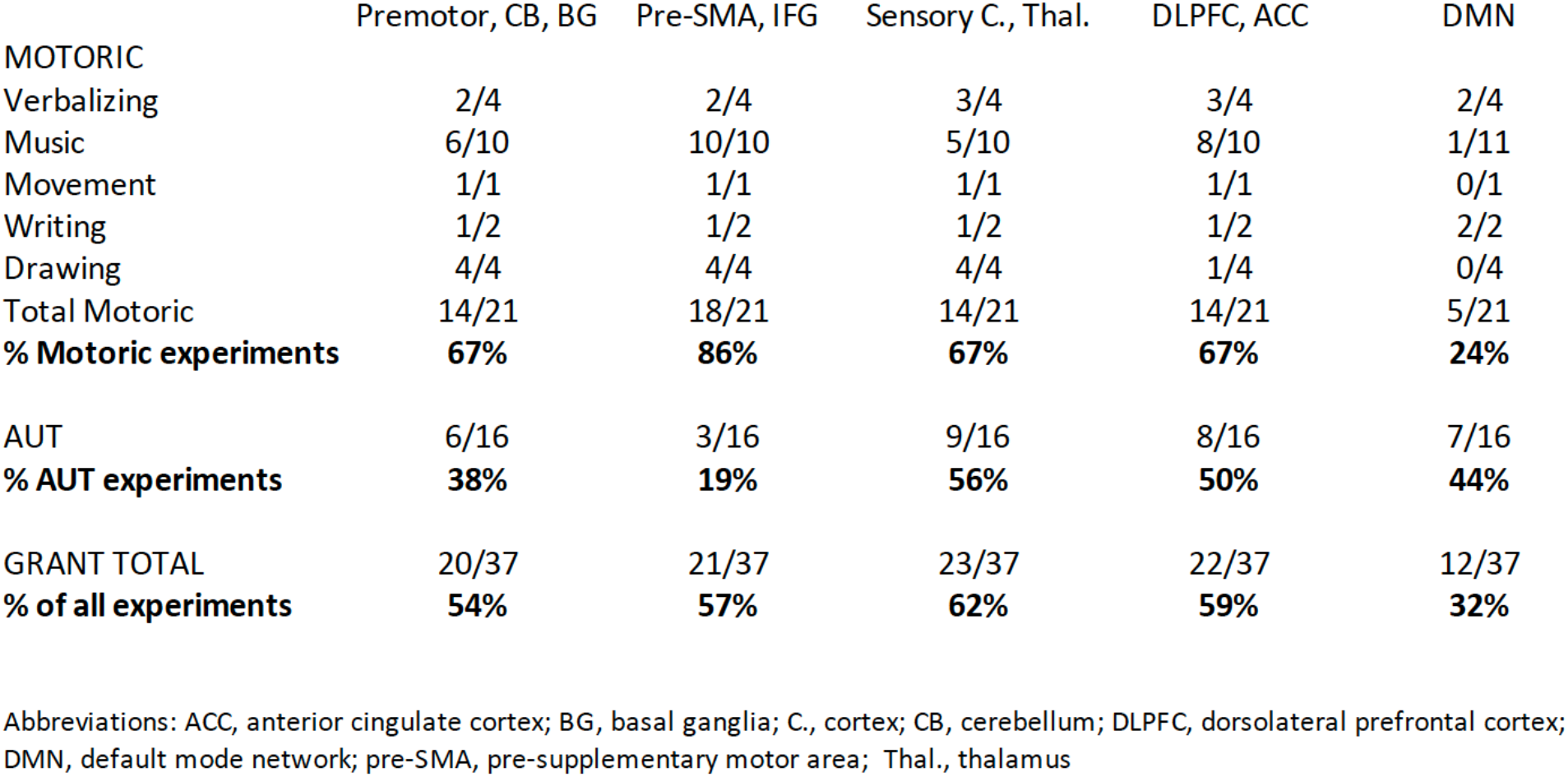
Contribution Analysis for the meta-analyses.

**Figure 5.**
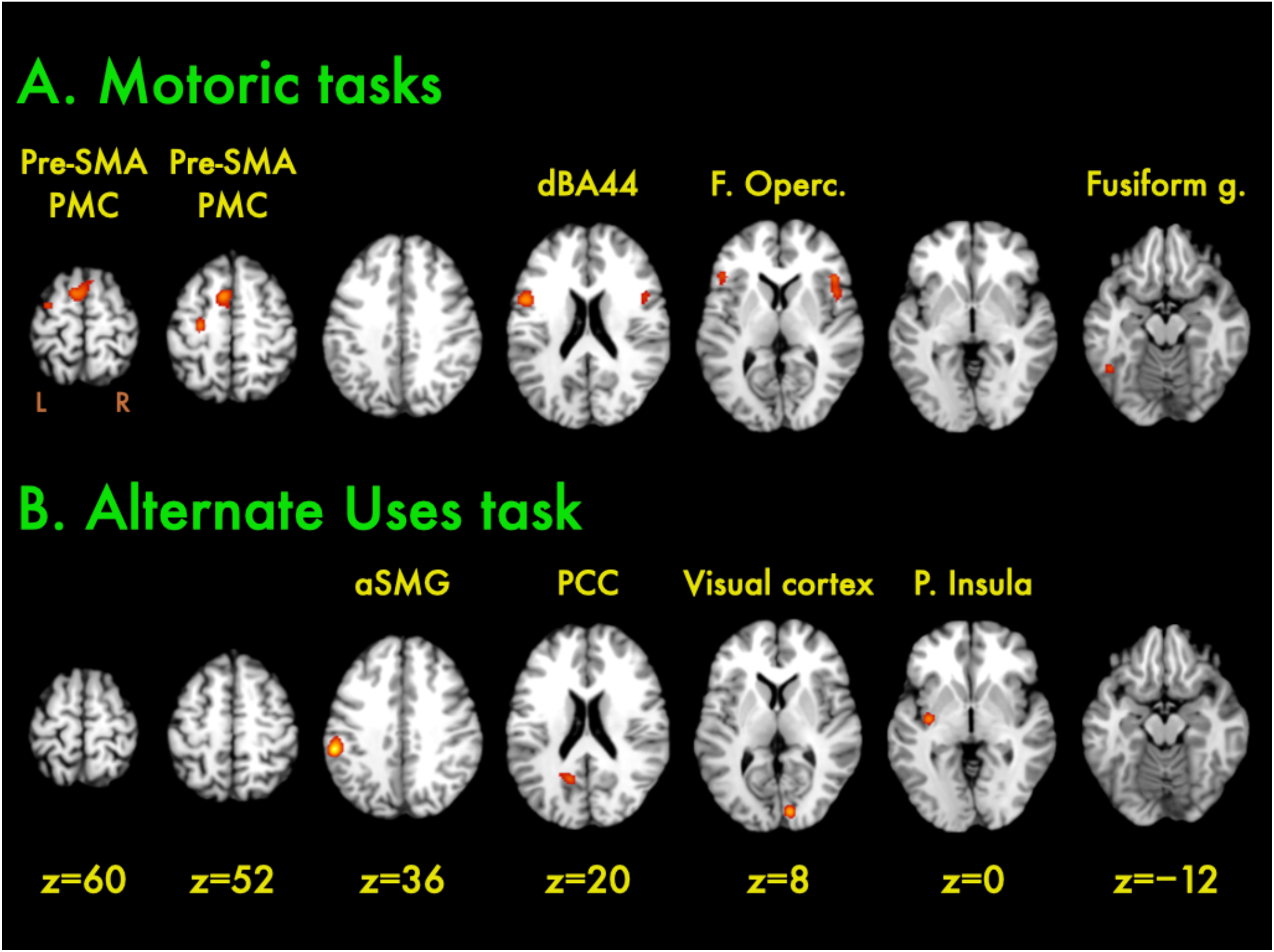
ALE clusters for the meta-analyses. Meta-analysis results for a) the motoric meta-analysis and b) the AUT analysis. The Talairach z level is indicated below each column of slices. The left side of the slice (L) is the left side of the brain. Abbreviations: aSMG, the anterior part of the supramarginal gyrus; dBA44, the dorsal part of Brodmann area 44; F. Operc., frontal operculum; Fusiform g., fusiform gyrus; PCC, posterior cingulate cortex; P. insula, posterior insula; PMC, premotor cortex; pre-SMA, pre-supplementary motor area.

Figure 5b shows the ALE foci for the AUT. The pattern shows no overlap with the motoric analysis. Not surprisingly, then, a conjunction analysis of the motoric and AUT maps yielded no significant clusters at p<0.05. The strongest ALE cluster for the AUT analysis was found in the anterior part of the supramarginal gyrus (aSMG) in left BA 40. Another pair of clusters was found in the left PCC, which was present in five of the 16 studies (31%). Additional clusters were found in the primary visual cortex (BA 17) and the posterior insula. Domain-general areas like the ACC, medial prefrontal cortex, and lateral prefrontal cortex were present in about half of the publications, but did not have sufficiently concordant locations to form ALE clusters. The CA shows a less reliable profile of activations across the 16 AUT papers than in the motoric meta-analysis. All of the categories of areas showed lower percentages of sourcing in the AUT analysis than in the motoric analysis (see Table 2). This is surprising given the fact that all of the AUT studies employed a single task, whereas the motoric analysis used a wide diversity of production tasks (e.g., jazz improvisation compared to creative drawing). Overall, the results for this divergent-thinking task were quite divergent.

## Discussion

We carried out two parallel meta-analyses of short-term creative processing, one a cross-modal analysis of motoric improvisation tasks and the other an analysis of the most universal divergent-thinking task used in the psychology literature, the Alternate Uses task. The two analyses presented different pictures of the neuroscience of creativity, with non-overlapping results at the standard thresholds used in ALE analyses (Eickhoff et al., 2016). The motoric meta-analysis produced highly reliable results across experiments, despite the fact that the studies used a wide diversity of creative tasks spanning five domains of production, including verbal improvisation, piano improvisation, movement improvisation, creative writing, and creative drawing, with manual production being the strongest link among them. In contrast to this, the AUT meta-analysis showed less-reliable results, despite employing a single task across all experiments. The conjunction analysis yielded no significant ALE clusters. Generalizations about the neural basis of creativity need to be tendered by this poor overlap between the networks for motoric creativity and divergent thinking, as well as by the relatively unreliable activation profile for studies using the AUT as the means for gauging creative cognition.

### Domain-specificity and -generality

Because all of the contrasts used in the meta-analyses contained a matched control condition that was designed to wash out sensorimotor activations associated with task production, we propose that the ALE results argue for a model of the neural basis of creativity that prioritizes domain-specific sensorimotor enhancement over domain-general expansion, as well as propose that motor-planning areas like the pre-SMA and IFG provide an ideal interface for uniting domain-specificity and -generality in short-term creative cognition. Creative processing seems to involve an enhancement of brain areas mediating non-creative or less-creative processing for that domain. This view might jibe with Weisberg’s (1993, 2006, 2020) perspective that creative cognition is a variant of ordinary thinking within a given domain, a process that he calls “thinking inside the box”. The involvement of sensorimotor areas in creative production has been underappreciated compared to other brain networks that have been implicated in creativity, such as the DMN. However, the results of the meta-analyses point to a clear role for sensorimotor areas in creativity, as seen not only in the ALE analyses themselves but in a majority of the contributing experiments in the CA. In fact, visual areas were the single most common type of area across the 37 studies in the meta-analyses.

Beyond sensorimotor areas, domain-general regions such as the DLPFC and ACC were present in many of the publications included in the meta-analyses, suggesting that creative production involves an expansion of cognitive resources beyond those used for non-creative production, perhaps involving increases in attention and/or working memory. However, these areas did not show sufficient spatial concordance across studies to appear as ALE clusters. Overall, if one looks across the individual studies included in the meta-analyses, one can find studies in which sensorimotor areas appear without the presence of domain-general areas (e.g., Fink et al., 2015; Sun et al., 2016). By contrast, there is no single study that reports the reverse, for example just the DLPFC, ACC, or PCC without the presence of motor and/or sensory areas as well. This observation would tend to prioritize sensorimotor areas over domain-general areas in terms of a hierarchy of involvement in creative cognition.

A number of the motoric articles have highlighted the role of deactivations in improvisation tasks when compared with motorically-matched control tasks (e.g., Donnay et al., 2014; Limb and Braun, 2008; Liu et al., 2015). We did not identify a sufficient number of experiments to permit us to meta-analyze deactivations. However, we note that the deactivations that are reported in the published studies occur overwhelmingly in *domain-general* brain areas, including DMN components such as the PCC/precuneus and medial prefrontal cortex, as well as in lateral parts of the prefrontal and parietal cortices. These are areas that are more active in the non-creative control tasks than in the creative tasks in these motoric studies. Whether this suggests that creativity is predicated on cortical disinhibition or not is in need of further exploration.

The most prominent areas of activation in the motoric meta-analysis were the left pre-SMA and bilateral IFG. As a group, these areas were present in 86% of the individual motoric experiments. They were also prominent in the meta-analyses of Boccia et al. (2015) and Chen et al. (2020). These areas are important for a neural model of creativity since they straddle the divide between domain-specificity and -generality (Marvel et al., 2019). On the one hand, they are premotor areas with connectivity to the primary motor cortex that play an undeniable role in motor planning and preparation. On the other hand, they are parts of domain-general networks, such as the cognitive control network (Cole and Schneider, 2007) and multiple drafts network (Duncan, 2010), that mediate domain-general functions such as working memory, action selection, and inhibition. Creativity – and complex behaviour more generally – depends on how domain-general processes impact sensorimotor mechanisms that are modally linked to task performance. In addition, premotor areas that are activated in a cross-modal manner, as the pre-SMA and IFG were in the motoric meta-analysis, have features that could be thought of as domain-general or at least that blur the dichotomy between domain-specific and -general.

Among the secondary motor areas that were observed in the motoric meta-analysis, the pre-SMA emerged as something of a cross-modal hot-spot for motoric improvisation, suggesting that it may serve as a potential hub in the brain’s improvisation network. Ruan et al. (2018) employed Meta-Analytic Connectivity Modeling to analyze the functional connections of the pre-SMA. These connections encompass many of the ALE clusters seen in the motoric meta-analysis, as well as non-ALE regions reported in the CA, including the IFG, premotor cortex, cerebellum (lobule VI), fusiform gyrus, inferior parietal lobule, putamen, and thalamus. Overall, not only is the pre-SMA the most widespread activation across all studies in the motoric meta-analysis, but the network for creative production maps quite well onto the zone of connectivity of the pre-SMA. In addition, while we did not observe the pre-SMA in our AUT analysis, it is reported in Cogdell-Brooke et al.’s (2020) meta-analysis of divergent thinking that includes other task-types beyond the AUT.

The pre-SMA, in addition to linking domain-specificity and -generality, might also provide an important linkage between creativity and *expertise*, since creative work in many domains, not least those of improvisation, requires expertise in a domain, where expertise can be conceptual as much as motoric. Along these lines, Pinho et al. (2014) found that functional connectivity with the pre-SMA increased with the total number of hours of improvisation training in classical and jazz pianists. Villarreal et al. (2013), in a study of rhythm improvisation, found greater levels of pre-SMA activation in their high-creativity group than their low-creativity group. de Aquino et al. (2019), in another study of rhythm improvisation, observed pre-SMA activation in both their musician and non-musician groups, but found relatively greater activity in the musicians; there was also greater activity in the left IFG. Dhakal et al. (2019) demonstrated that the pre-SMA’s role in vocal musical improvisation occurred not just during motor production but during a condition of mental imagery as well. Finally, Bashwiner et al. (2016), in structural study of musical expertise, found that the cortical surface area of the pre-SMA was associated with increasing ratings of personal creativity, as measured by means of self-reports of experience at improvisation and composition.

These findings about the functions of the pre-SMA are reinforced by anatomical connectivity as well. The premotor areas seen in the motoric analysis are interconnected via a diagonal fasciculus known as the *frontal aslant tract* (FAT) that connects the inferior frontal gyrus with the SMA/pre-SMA in the medial part of the superior frontal gyrus (Briggs et al., 2019; Catani et al., 2012; Dick et al., 2019; Ford et al., 2010). The FAT has been implicated in higher-order cognition, including language functioning and working memory (Dick et al., 2019). It has also been implicated in fluid intelligence (P. Y. Chen et al., 2020), which is seen as being a critical factor in creative cognition (Vartanian, 2019, 2013). The results of the current meta-analysis open the door to examining the structural properties of the FAT in diffusion-imaging analyses of the white-matter correlates of creativity, both cross-sectionally and longitudinally. For example, highly creative people might differ from less-creative people in the fractional isotropy or mean diffusivity of the FAT.

### Divergent thinking

While the AUT is typically regarded as a purely ideational task, we propose interpreting its fMRI activation profile in a more modal manner. After all, the AUT was considered as a modal task by its creators, where it was classified as a “verbal” form of DT, as compared to “figural” tasks (Guilford, 1967; Torrance, 1974). The most prominent ALE cluster in the AUT meta-analysis, as well as in the DT meta-analyses of Wu et al. (2015) and Cogdell-Brooke et al. (2020), was located in the anterior part of the supramarginal gyrus (aSMG) in left BA 40 (see also Gonen-Yaacovi, 2013). Cogdell-Brooke et al. (2020) argued that this region is part of a network for tool manipulation, and we would like to expand on their proposal. The ALE cluster for the aSMG – at Talairach coordinate -54, -32, 36 – is quite proximate to the left inferior parietal area that Orban and Caruana (2014) have implicated in both tool use and the observation of tool use and hand actions, hence a region of sensorimotor overlap. It is an area that Orban (2016) has referred to as “the tool-use area”, a region showing left-hemisphere asymmetry. The AUT can be thought of as a covert objection-manipulation task employing visual and motor imagery. This contention is supported by an ALE cluster in the primary visual cortex (BA 17). While the posterior insula’s role in the AUT is not clear, Kurth et al.’s (2010) large-scale ALE meta-analysis of the insula linked the posterior insula primarily with sensory and motor functioning. Gharizi et al.’s (2017) diffusion-based structural connectivity analysis of the human insula found prominent connections between the posterior insula and the supramarginal gyrus as well as the posterior part of the cingulate cortex and perhaps visual cortex. The ALE regions of the AUT meta-analysis seem to be connected with one another both functionally and structurally. Therefore, instead of conceiving of the AUT as “the generation of novel ideas”, the results of the meta-analysis point to a much more domain-specific interpretation of the AUT as being related to an enhancement of the visual/motor imagery of object manipulation, not least since all of the objects used in the AUT are manipulable objects. Future work on the AUT should take a more modal approach to this task. Matheson and Kenett (2020) have provided an action-simulation account of the AUT that takes important steps in this direction. As they point out, “simulations of actions (and tool-related action in particular) support generating creative uses of objects” when performing the AUT (p. 2).

The AUT has been strongly linked with the DMN in both functional-connectivity analyses (Beaty et al., 2018, 2015, 2014) and structural analyses of DT (Jung et al., 2013; Kühn et al., 2014; Wertz et al., 2020), although far less so in standard functional-activation studies. While neither of the two previous DT meta-analyses reported DMN components in their results (Cogdell-Brooke et al., 2020; Wu et al., 2015), we observed a PCC cluster in the AUT analysis in left BA 23/31 close to where Beaty et al. (2015) reported their most extensive cluster in a multivoxel pattern analysis of the AUT. The contribution analysis revealed that, with one exception (Benedek et al., 2018), all of the AUT studies that showed PCC activations were those in which participants performed the task covertly, rather than reporting their ideas vocally during the task. (In fact, 10 of the 16 AUT studies in the meta-analysis used button press to register idea occurrence, rather than a vocal reporting of the alternate uses.) Given the fact that the DMN is well-known to favor internal processing over external processing of information (Beaty et al., 2015), the PCC activation might relate to this internal performance of the task. For the motoric ALE, the PCC was present in only one of the 21 contributing studies. This observation would tend to limit the generalizability of DMN findings coming from studies of the AUT to other domains and tasks of creativity. In addition, the AUT studies showed less reliability of findings than did the motoric studies, with more similarity of results being found within labs (e.g., the studies of Fink et al., 2015, 2010, 2009 and those of Abraham et al., 2018, 2012) than among labs. Activations in visual areas were the most common link across all of the AUT studies. Such areas have been implicated in structural studies of divergent thinking (Jung et al., 2010).

It is unclear if the unique presence of the PCC in the AUT (although in only 5 of 16 experiments) is related to the generative nature of the task, as compared to the more elaborative nature of the motoric tasks. There have been proposals that the DMN is the principal generative source of creative ideas (Beaty et al., 2016; Jung et al., 2013). However, Ellamil et al.’s (2012) study of creative drawing found the PCC to be preferentially associated with the evaluative phase of their task, not the generative phase. Liu et al.’s (2015) study of poetry improvisation found the PCC to be more active during the replicative control task (i.e., typing out pre-learned poems) than in the creative generation task. Erhard et al. (2014) reported no difference between the generative brainstorming phase of creative writing and a passive reading task. Finally, as just mentioned, more than half of the AUT studies that were used in the meta-analysis did not show DMN components in their creative-vs.-non-creative contrasts. Based on the results of the present meta-analyses, it is premature to assign brain areas to the generation vs. elaboration phases of creative production. This should be an important goal of future studies that look specifically at multi-phase tasks.

### Comparison with other ALE meta-analyses

The current meta-analysis is a first attempt to compare motor improvisation with divergent thinking in a directed manner through separate analyses and then a conjunction analysis. It shares a number of the contributing experiments with the previously published ALE meta-analyses of creativity, and so it is not surprising that the results show similarities with them. Regarding the motoric analysis, we limited our inclusion to experiments that had a matched creative-vs.-non-creative contrast, whereas the previous studies also included experiments that had contrasts against rest or fixation, which would lead to more-extensive activation profiles than using a control condition matched for sensorimotor demands. Despite this more conversative approach, our results ended up being quite similar to the motoric results reported by Boccia et al. (2015) and Chen et al. (2020), with an emphasis on the pre-SMA, IFG, and PMC.

Regarding the AUT analysis, previous meta-analyses of divergent thinking have included other tasks beyond the AUT, for example verb generation, visual imagery, metaphor production, and creative drawing (Cogdell-Brooke et al., 2020; Wu et al., 2015). As a result, key differences were observed between the current meta-analysis and the previous two. In particular, neither of the previous analyses reported ALE clusters in the PCC, visual cortex, or posterior insula, all of which were present in at least one third of the AUT studies. So, it is possible that the inclusion of additional task-types in the other meta-analyses diluted the contribution of these three areas to the ALE results. This is important to keep in mind given that the PCC has been a focal point of connectivity-based analyses of DT (Beaty et al., 2018, 2015). On the other hand, Gonen-Yaacovi et al. (2013) and Cogdell-Brooke et al. (2020) reported significant ALE clusters in the pre-SMA and frontal operculum that were absent in our AUT meta-analysis, arguing that these clusters most likely originated in the non-AUT experiments. For example, verb generation is a common activator of the frontal operculum (e.g., Warburton et al., 1996). Our conjunction analysis would most likely have contained the pre-SMA and frontal operculum had we included other categories of tasks in our DT analysis. The contribution analysis for the AUT revealed that the pre-SMA was present in only a single study (Abraham et al., 2018) and the frontal operculum in only two studies (Abraham et al., 2018, 2012), all from the same lab group.

### Future prospects: Short-term phases and long-term creativity

For the neuroscience of creativity to advance, it needs to broach the two overarching dimensions of domain and time-frame that were discussed in the introductory sections (see Figure 2). For the former, this means looking at creativity in a cross-modal manner, as attempted here and in the previously published meta-analyses. For the latter, it involves addressing the iterative and explorational nature of creative cognition, as well as making initial steps toward looking at long-term creativity. The AUT has been a dominant task in the creativity field (Benedek et al., 2019), but it is restricted to a generative phase and lacks any sense of elaboration or revision. It contrasts with all of the tasks used in the motoric meta-analysis that not only require idea generation but also a great deal of elaboration over time to either maintain continuity in performance (as in jazz improvisation) or to flesh out an idea to generate a short-term product (as in creative drawing). As was discussed in the introductory sections, real-world creativity is not based purely or even prominently on brainstorming alone, but requires extensive exploration and revision to test out the validity and workability of ideas (Cropley, 2006; Weisberg, 2020). The psychology of creativity has been highly idea-centric and individualist since its origins. However, there is nothing magical about idea generation. It is merely one part of a long and arduous process of doing creative work. While brainstorming might be a contributor to idea generation, the real work of creativity is in the exploration and validation of these ideas (Wallas, 1926/2014; Finke et al., 1992) as well in coping with the critical reception of consumers, whose actions determine whether the creative product is considered valuable or not.

Long-term creativity will be an important frontier for future work on the neuroscience of creativity. While long-term creativity has been looked at in ethnographic psychological studies (e.g., Mace and Ward, 2002; Ness and Dysthe, 2020), it has not be examined in neural studies to date, and there are clear analytical challenges to doing so. A great deal of methodological creativity will be required to examine the neural basis of long-term creative processing, but that outcome should be seen as being one of the ultimate goals of the field. When it comes to studies of short-term creativity, only a small number of studies have examined the phases of creativity in order to look at processes like revision and evaluation. Liu et al. (2015) had both trained poets and novices improvise poems (through typing) during a generation phase and then revise the poems during a second phase. The pre-SMA and frontal operculum were present in the generation phase (which itself involves much elaboration), whereas the dorsal part of BA 44 was present in revision, but not generation. A similar finding was observed in Ellamil et al.’s (2012) study of creative drawing, in which the generation of a drawing was compared with a later evaluation. Dorsal BA 44 was present in the “evaluate > generate” contrast. However, so was the frontal operculum, which was preferentially present in the generation phase in Liu et al. (2015). Hence, the limited information that exists from multi-phase studies of creativity suggests that BA 44 might double-duty between generation and exploration/revision/evaluation processes. The dorsal part of BA 44 might have a more prominent role in exploration, rather than generation. Overall, future studies need to address not just the one-shot generation of ideas but the elaboration and exploration of such ideas, which is the critical part of creativity in virtually all domains.

Finally, the neuroscience of real-world creativity is going to have to address the overwhelmingly collaborative nature of creative work (Fischer et al., 2005; John-Steiner, 2000; Kimmel et al., 2018; Ness and Dysthe, 2020; Sawyer, 2007, 2019; Sawyer and DeZutter, 2009), for example the collaborative scientific work that underlies illuminating the neural basis of creativity. There is no question that the future of the field will one day be dominated by work on collaborative creativity as well as its social determinants and effects. Creativity and innovation are intimately linked with the mechanisms of the cultural evolution of products (Brown, 2021; Gabora, 2019). Thus far, there are a number of promising analyses of collaborative creativity using fMRI (Chauvigné et al., 2018; Xie et al., 2020) and functional near infrared spectroscopy (Lu et al., 2019). The analysis of collaborative creativity will require an understanding not only of individual-level creativity, but of the mechanisms of partnering, joint action, and social interaction as well (Gallotti et al., 2017; Pacherie, 2011; Redcay and Schilbach, 2019; Sebanz et al., 2006) and how these mechanisms influence creative cognition. This impacts not just the generation of ideas, but their evaluation, exploration, elaboration, revision, and ultimately how the critical reception of creative products by consumers feeds back to influence the ideas of creators. Creators are embedded not only within domains but within broad cultural systems that interact with creators bidirectionally (Csikszentmihalyi, 1988; Glǎveanu, 2010), serving both as a source of inspiration for creative work and as the intended recipients for one’s creative output.

## Conclusions

We carried out two parallel meta-analyses of short-term creativity, comparing motoric creativity (incorporating studies from five domains of improvisation) with the standard laboratory test of divergent thinking (the AUT). All of the experiments that were included in the meta-analyses contained a contrast between a creative task and a matched non-creative or less-creative control task in order to equate the sensorimotor demands of task performance. We argue that the results of both meta-analyses can be interpreted as being primarily driven by sensorimotor effects, where creative processing is an enhancement of brain areas mediating non-creative or less-creative processing for that domain. Domain-general areas are critical as well, but seem to occupy a lower tier in the hierarchy. An interface between domain-specificity and -generality was found in the pre-SMA, dorsal IFG, and frontal operculum, which might unite motor planning and executive control. These areas are interconnected via the frontal aslant tract, suggesting that this tract might be a useful target in white-matter analyses of creativity. Future work on the neuroscience of creativity needs to move beyond one-shot brainstorming tasks to look at multi-phase tasks that involve elaboration and exploration in addition to generation. Ultimately, the creativity field needs to devise creative approaches to studying long-term explorational creativity, the kind that underlies scientific discovery, technology development, and artistic creation. This needs to include the important role of collaborative creativity that is prominent in virtually all domains of creative work.

## Acknowledgments

This work was supported by a grant to S.B. from Natural Sciences and Engineering Research Council (NSERC) of Canada (grant number RGPIN-2020-05718). We are grateful to Simon Eickhoff for advice on the analysis and to Zoe Lazar-Kurz for critical comments on the manuscript.

## Notes

### Competing Interest Statement

The authors have declared no competing interest.

