## Supplementary Table 1 for "The neural basis of creative production: A cross-modal ALE meta-analysis"

**Table S1: Reference list of the articles contributing to the meta-analysis, organized by category**


VERBALIZATION

Amir, O., Biederman, I., Graham, D.J., Meng, M., Brownell, H.H., 2016. The neural correlates of humor creativity. Front. Hum. Neurosci. 10, 597. https://doi.org/10.3389/fnhum.2016.00597

Bechtereva, N.P., Korotkov, A.D., Pakhomov, S. V., Roudas, M.S., Starchenko, M.G., Medvedev, S. V., 2004. PET study of brain maintenance of verbal creative activity. Int. J. Psychophysiol. 53, 11–20. https://doi.org/10.1016/j.ijpsycho.2004.01.001

Howard-Jones, P.A., Blakemore, S.J., Samuel, E.A., Summers, I.R., Claxton, G., 2005. Semantic divergence and creative story generation: An fMRI investigation. Cogn. Brain Res. 25, 240–250. https://doi.org/10.1016/j.cogbrainres.2005.05.013

Liu, S., Chow, H.M., Xu, Y., Erkkinen, M.G., Swett, K.E., Eagle, M.W., Rizik-Baer, D.A., Braun, A.R., 2012. Neural correlates of lyrical improvisation: An fMRI study of freestyle rap. Sci. Rep. 2, 834. https://doi.org/10.1038/srep00834

MUSIC

Bengtsson, S.L., Csíkszentmihályi, M., Ullén, F., 2007. Cortical regions involved in the generation of musical structures during improvisation in pianists. J. Cogn. Neurosci. 19, 830–842. https://doi.org/10.1162/jocn.2007.19.5.830

Berkowitz, A.L., Ansari, D., 2008. Generation of novel motor sequences: The neural correlates of musical improvisation. Neuroimage 41, 535–543. https://doi.org/10.1016/j.neuroimage.2008.02.028

de Aquino, M.P.B., Verdejo-Román, J., Pérez-García, M., Pérez-García, P., 2019. Different role of the supplementary motor area and the insula between musicians and non-musicians in a controlled musical creativity task. Sci. Rep. 9, 13006. https://doi.org/10.1038/s41598-019-49405-5

de Manzano, Ö., Ullén, F., 2012a. Activation and connectivity patterns of the presupplementary and dorsal premotor areas during free improvisation of melodies and rhythms. Neuroimage 63, 272–280. https://doi.org/10.1016/j.neuroimage.2012.06.024

de Manzano, Ö., Ullén, F., 2012b. Goal-independent mechanisms for free response generation: Creative and pseudo-random performance share neural substrates. Neuroimage 59, 772–780. https://doi.org/10.1016/j.neuroimage.2011.07.016

Dhakal, K., Norgaard, M., Adhikari, B.M., Yun, K.S., Dhamala, M., 2019. Higher node activity with less functional connectivity during musical improvisation. Brain Connect. 9, 296–309. https://doi.org/10.1089/brain.2017.0566

Donnay, G.F., Rankin, S.K., Lopez-Gonzalez, M., Jiradejvong, P., Limb, C.J., 2014. Neural substrates of interactive musical improvisation: An fMRI study of “trading fours” in jazz. PLoS One 9, e88665. https://doi.org/10.1371/journal.pone.0088665

Limb, C.J., Braun, A.R., 2008. Neural substrates of spontaneous musical performance: An fMRI study of jazz improvisation. PLoS One 3, e1679. https://doi.org/10.1371/journal.pone.0001679

McPherson, M.J., Barrett, F.S., Lopez-Gonzalez, M., Jiradejvong, P., Limb, C.J., 2016. Emotional intent modulates the neural substrates of creativity: An fMRI study of emotionally targeted improvisation in jazz musicians. Sci. Rep. 6, 18460. https://doi.org/10.1038/srep18460

Villarreal, M.F., Cerquetti, D., Caruso, S., Schwarcz López Aranguren, V., Gerschcovich, E.R., Frega, A.L., Leiguarda, R.C., 2013. Neural correlates of musical creativity: Differences between high and low creative subjects. PLoS One 8, e75427. https://doi.org/10.1371/journal.pone.0075427

<u>MOVEMENT</u>

Chauvigné, L.A.S., Belyk, M., Brown, S., 2018. Taking two to tango: fMRI analysis of improvised joint action with physical contact. PLoS One 13, e0191098. https://doi.org/10.1371/journal.pone.0191098

<u>WRITING</u>

Liu, S., Erkkinen, M.G., Healey, M.L., Xu, Y., Swett, K.E., Chow, H.M., Braun, A.R., 2015. Brain activity and connectivity during poetry composition: Toward a multidimensional model of the creative process. Hum. Brain Mapp. 36, 3351–3372. https://doi.org/10.1002/hbm.22849

Shah, C., Erhard, K., Ortheil, H.J., Kaza, E., Kessler, C., Lotze, M., 2013. Neural correlates of creative writing: An fMRI Study. Hum. Brain Mapp. 34, 1088–1101. https://doi.org/10.1002/hbm.21493

<u>DRAWING</u>

Ellamil, M., Dobson, C., Beeman, M., Christoff, K., 2012. Evaluative and generative modes of thought during the creative process. Neuroimage 59, 1783–1794. https://doi.org/10.1016/j.neuroimage.2011.08.008

Hahm, J., Kim, K.K., Park, S.-H., Lee, H.-M., 2017. Brain areas subserving Torrance Tests of Creative Thinking: An functional magnetic resonance imaging study. Dement. Neurocognitive Disord. 16, 48–53. https://doi.org/10.12779/dnd.2017.16.2.48

Park, H.R.P., Kirk, I.J., Waldie, K.E., 2015. Neural correlates of creative thinking and schizotypy. Neuropsychologia 73, 94–107. https://doi.org/10.1016/j.neuropsychologia.2015.05.007

Saggar, M., Quintin, E.M., Kienitz, E., Bott, N.T., Sun, Z., Hong, W.C., Chien, Y.H., Liu, N., Dougherty, R.F., Royalty, A., Hawthorne, G., Reiss, A.L., 2015. Pictionary-based fMRI paradigm to study the neural correlates of spontaneous improvisation and figural creativity. Sci. Rep. 5, 1–11. https://doi.org/10.1038/srep10894

<u>ALTERNATE USES TASK</u>

Abdul Hamid, K., Yusoff, A.N., Rahman, S., Osman, S.S., Azmi, N.H., Surat, S., Ahmad Marzuki, M., 2019. Cortical responses during divergent thinking tasks after creativity stimulation. Psychol. Neurosci. 12, 342–362. https://doi.org/10.1037/pne0000168

Abraham, A., Pieritz, K., Thybusch, K., Rutter, B., Kröger, S., Schweckendiek, J., Stark, R.,

Windmann, S., Hermann, C., 2012. Creativity and the brain: Uncovering the neural signature of conceptual expansion. Neuropsychologia 50, 1906–1917. https://doi.org/10.1016/j.neuropsychologia.2012.04.015

Abraham, A., Rutter, B., Bantin, T., Hermann, C., 2018. Creative conceptual expansion: A combined fMRI replication and extension study to examine individual differences in creativity. Neuropsychologia 118, 29–39. https://doi.org/10.1016/j.neuropsychologia.2018.05.004

Benedek, M., Schües, T., Beaty, R.E., Jauk, E., Koschutnig, K., Fink, A., Neubauer, A.C., 2018. To create or to recall original ideas: Brain processes associated with the imagination of novel object uses. Cortex 99, 93–102. https://doi.org/10.1016/j.cortex.2017.10.024

Fink, A., Benedek, M., Koschutnig, K., Pirker, E., Berger, E., Meister, S., Neubauer, A.C., Papousek, I., Weiss, E.M., 2015. Training of verbal creativity modulates brain activity in regions associated with language- and memory-related demands. Hum. Brain Mapp. 36, 4104–4115. https://doi.org/10.1002/hbm.22901

Fink, A., Grabner, R.H., Benedek, M., Reishofer, G., Hauswirth, V., Fally, M., Neuper, C., Ebner, F., Neubauer, A.C., 2009. The creative brain: Investigation of brain activity during creative problem solving by means of EEG and fMRI. Hum. Brain Mapp. 30, 734–748. https://doi.org/10.1002/hbm.20538

Fink, A., Grabner, R.H., Gebauer, D., Reishofer, G., Koschutnig, K., Ebner, F., 2010. Enhancing creativity by means of cognitive stimulation: Evidence from an fMRI study. Neuroimage 52, 1687–1695. https://doi.org/10.1016/j.neuroimage.2010.05.072

Heinonen, J., Numminen, J., Hlushchuk, Y., Antell, H., Taatila, V., Suomala, J., 2016. Default mode and executive networks areas: Association with the serial order in divergent thinking. PLoS One 11, e0162234. https://doi.org/10.1371/journal.pone.0162234

Ivancovsky, T., Kleinmintz, O., Lee, J., Kurman, J., Shamay-Tsoory, S.G., 2018. The neural underpinnings of cross-cultural differences in creativity. Hum. Brain Mapp. 39, 4493–4508. https://doi.org/10.1002/hbm.24288

Madore, K.P., Thakral, P.P., Beaty, R.E., Addis, D.R., Schacter, D.L., 2019. Neural mechanisms of episodic retrieval support divergent creative thinking. Cereb. Cortex 29, 150–166. https://doi.org/10.1093/cercor/bhx312

Mayseless, N., Eran, A., Shamay-Tsoory, S.G., 2015. Generating original ideas: The neural underpinning of originality. Neuroimage 116, 232–239. https://doi.org/10.1016/j.neuroimage.2015.05.030

Sun, J., Shi, L., Chen, Q., Yang, W., Wei, D., Zhang, J., Zhang, Q., Qiu, J., 2019. Openness to experience and psychophysiological interaction patterns during divergent thinking. Brain Imaging Behav. 13, 1580–1589. https://doi.org/10.1007/s11682-018-9965-2

Vartanian, O., Beatty, E.L., Smith, I., Blackler, K., Lam, Q., Forbes, S., 2018. One-way traffic: The inferior frontal gyrus controls brain activation in the middle temporal gyrus and inferior parietal lobule during divergent thinking. Neuropsychologia 118, 68–78. https://doi.org/10.1016/j.neuropsychologia.2018.02.024

Vartanian, O., Bouak, F., Caldwell, J.L., Cheung, B., Cupchik, G., Jobidon, M.E., Lam, Q., Nakashima, A., Paul, M., Peng, H., Silvia, P.J., Smith, I., 2014. The effects of a single night of sleep deprivation on fluency and prefrontal cortex function during divergent thinking. Front. Hum. Neurosci. 8, 214. https://doi.org/10.3389/fnhum.2014.00214
